# Fixel-Based Analysis and Free Water Corrected DTI Evaluation of HIV Associated Neurocognitive Disorders

**DOI:** 10.1101/2021.05.06.443022

**Authors:** Alan Finkelstein, Abrar Faiyaz, Miriam T. Weber, Xing Qiu, Md Nasir Uddin, Jianhui Zhong, Giovanni Schifitto

## Abstract

White matter damage is a consistent finding in HIV infected (HIV+) individuals. Previous studies have evaluated WM fiber tract specific brain regions in HIV-associated neurocognitive disorders using diffusion tensor imaging (DTI). However, DTI might lack an accurate biological interpretation, and the technique suffers from several limitations. Here, we sought to evaluate Fixel-based analysis (FBA) and free water corrected DTI (fwcDTI) metrics between HIV+ and HIV uninfected (HIV−) individuals, and their relationships with blood markers and cognitive scores. We also compared the specificity of both MRI metrics in their ability to distinguish between individuals with and without cognitive impairment using machine learning classifiers. Using 94 age-matched participants, we found that whole brain FBA was significantly reduced (up to 15%) in various fiber bundles. Tract based special statistics (TBSS) of fwcDTI metrics revealed decreased fractional anisotropy FA_T_ (by 1-2%) in HIV+ compared to HIV− individuals in areas consistent with those observed in FBA, but these were not significant. An adaptive boosting classifier reliably distinguished between cognitively normal patients and those with cognitive impairment with 80% precision and 78% recall. Therefore, FBA may serve as a potential in-vivo biomarker for evaluating and monitoring axonal degeneration in HIV+ patients at risk for neurocognitive impairment.

## 1 Introduction

Combined antiretroviral therapy (cART) has reduced morbidity and mortality rates significantly in HIV infected (HIV+) individuals (1). However, the increased survival may be masking an increase in cognitive impairment (2), mediated by injury to the central nervous system (CNS) and disruption of the blood brain barrier (BBB) (3). The HIV reservoir in the CNS resides primarily in microglia and perivascular macrophages, resulting in chronic neuroinflammation (4). While the small pool of infected cells in the CNS can release neurotoxic viral proteins, Tat and gp120, the larger pool of activated glia cells is responsible for the release of cytokines which can induce neuronal injury and cell death (5). HIV associated oligodendrocyte injury results in demyelination and alterations in white matter (WM) structural integrity (6). Thus, damage to WM fibers is likely a key factor in cognitive impairment observed in HIV associated neurocognitive disorder (HAND) (7).

MR neuroimaging studies have sought to identify potential *in vivo* biomarkers to investigate CNS injury in the setting of HIV infection (8). Structural and functional MRI have helped elucidate how atrophy and aberrant network topology mediate cognitive decline in HIV infection (9, 10). Nonetheless, given the well-established presence of WM alterations in HIV infection, it is paramount to further characterize WM in HAND. However, during chronic neuroinflammation, there may be contributing vasogenic edema (11), confounding the interpretation of WM lesions. Accordingly, appropriate models that accurately account for free water (FW) contamination are necessary to sufficiently evaluate WM structural integrity in HIV infection (12).

Due to its noninvasiveness, diffusion tensor imaging (DTI) has been widely used in clinical neuroimaging studies (13, 14). DTI metrics such as fractional anisotropy (FA), axial diffusivity (AD), radial diffusivity (RD) and mean diffusivity (MD) characterize the orientation and distribution of the random movements of water molecules, diffusion magnitude, diffusional directionality perpendicular to the axon, and diffusional directionality along the axon, respectively (15, 16). Previous studies have shown that FA is decreased in the posterior limb of internal capsule (PLIC), the corticospinal tract (CST), temporal and frontoparietal WM regions, whereas RD and MD were increased in bilateral CST, temporal and frontal WM regions in HIV-infected (HIV+) individuals (17–22). Decreased FA has also been observed in the superior longitudinal fasciculus (SLF) and was correlated with decreased memory and executive function in HIV+ subjects exhibiting HAND (23). Tract-based spatial statistics (TBSS) is a popular voxel-based method to analyze DTI metrics, which maps control and disease cohort FA images to a WM skeleton to improve correspondence between subjects (24). We have previously reported diffuse FA and MD abnormalities using TBSS in HIV+ individuals (25).

However, voxel based measures are often contaminated by extracellular free water (FW) (12). FW contamination in the diffusion signal is due to water molecules that are not restricted by their environment, such as the cerebrospinal fluid (CSF). Edema caused by stroke (26), brain tumors (12), or neuroinflammation can also contaminate WM voxels (27). Accordingly, FW contaminated voxels will fit more towards an isotropic tensor and exhibit decreased FA values, confounding the interpretation of the results. Previously, several studies have reported that FW correction enhances specificity of DTI metrics (28–31) and reduces test-retest reproducibility errors (32). However, the diffusion tensor model is limited in that it is not able to reliably model complex and crossing-fiber populations, which are present in up to 90% of WM voxels (33, 34). Furthermore, while TBSS of FW corrected FA (FA_T_) is likely to provide more reliable measures of WM integrity in HIV infection, it does not include orientation information.

Fixel-based analysis (FBA) is a recent technique that models individual fibers at the sub-voxel level, termed fixels, which allow tract-specific comparisons (35). FBA enables the characterization of multiple fiber populations within a voxel, circumventing interpretation issues that commonly arise with voxel-averaged measurements such as FA and MD. Moreover, FBA accounts for both macrostructural (fiber bundle) and microstructural (within voxel) changes within WM, providing a more comprehensive understanding of intra-axonal and fiber tract changes. Accordingly, FBA has been used in several neurological disorders including Parkinson’s disease (36, 37), multiple sclerosis (38, 39), traumatic brain injury (40, 41), schizophrenia (41) and healthy aging (42). FBA can be used to estimate fiber density (FD) within a fiber bundle, the fiber bundle cross section (FC) or a combined measure, fiber density cross section (FDC). FD is related to the intra-axonal volume, and a corresponding decrease in FD may reflect axonal degeneration (35). FD or the apparent fiber density (AFD) is calculated from the fiber orientation distribution (FOD) which is estimated using multi tissue constrained spherical deconvolution (MT-CSD) (43). By modeling grey matter (GM), WM, and CSF separately, MT-CSD accounts for FW contamination and has been shown to have better test-retest reliability than traditional DTI metrics (44). FC reflects the cross-sectional area of a fiber bundle, perpendicular to the length axis, and is derived from the Jacobian of the non-linear transformation from subject space to template space. Decreases in FC may result from WM atrophy and loss of the extra-axonal space, while increases may reflect inflammation and an increase in inflammatory proteins (35). FDC accounts for both macroscopic and microscopic effects on fiber density.

The aim of this study was to refine our understanding of how WM structural integrity is affected in HIV-infected individuals, and if these changes were associated with cognitive performance in HAND, using two approaches, FBA and fwcDTI. Additionally, we investigated whether fiber tract degeneration was related to inflammatory blood markers NfL and Tau of HIV infection. Machine learning classification, using a set of binary classifiers was also performed to distinguish cognitively normal individuals from those with cognitive impairment in HIV+ individuals.

## 2 Materials and Methods

### 2.1 Study Participants

Forty-two treatment-nai◻ve HIV+ participants (4 females and 40 males; mean age ± standard error, SE=34.48± 1.95 years, range 20-63 years) and 52 age-matched HIV uninfected (HIV−) participants (26 females and 26 males; mean age ± SE=37.02 ± 1.66 years, range 18-63 years) were enrolled in a study assessing the potential neurotoxicity of combination antiretroviral therapy treatment (cART) study at the University of Rochester Medical Center. All participants provided written informed consent before enrollment according to the institutional protocol and underwent clinical, laboratory and brain MRI exams. All experiments were performed in accordance with relevant guidelines and regulations. The data reported here reflect the baseline assessment of HIV+, cART naive individuals prior to starting cART. This time-point was chosen because it represents the clearest difference in cognitive performance within the HIV+ group. Details about study participants (including age, sex, clinical results) are provided in **Table 1**.

**Table 1.**
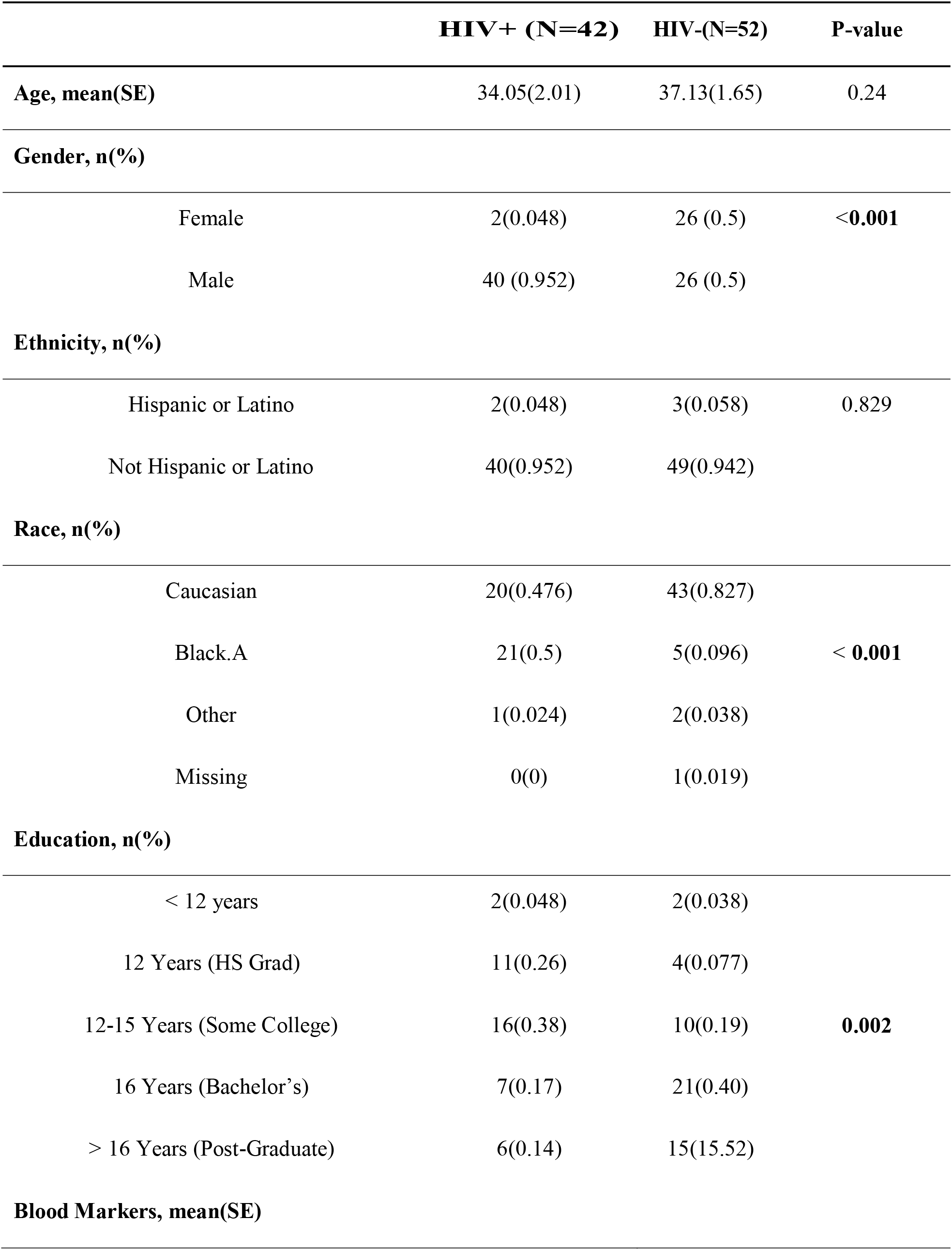
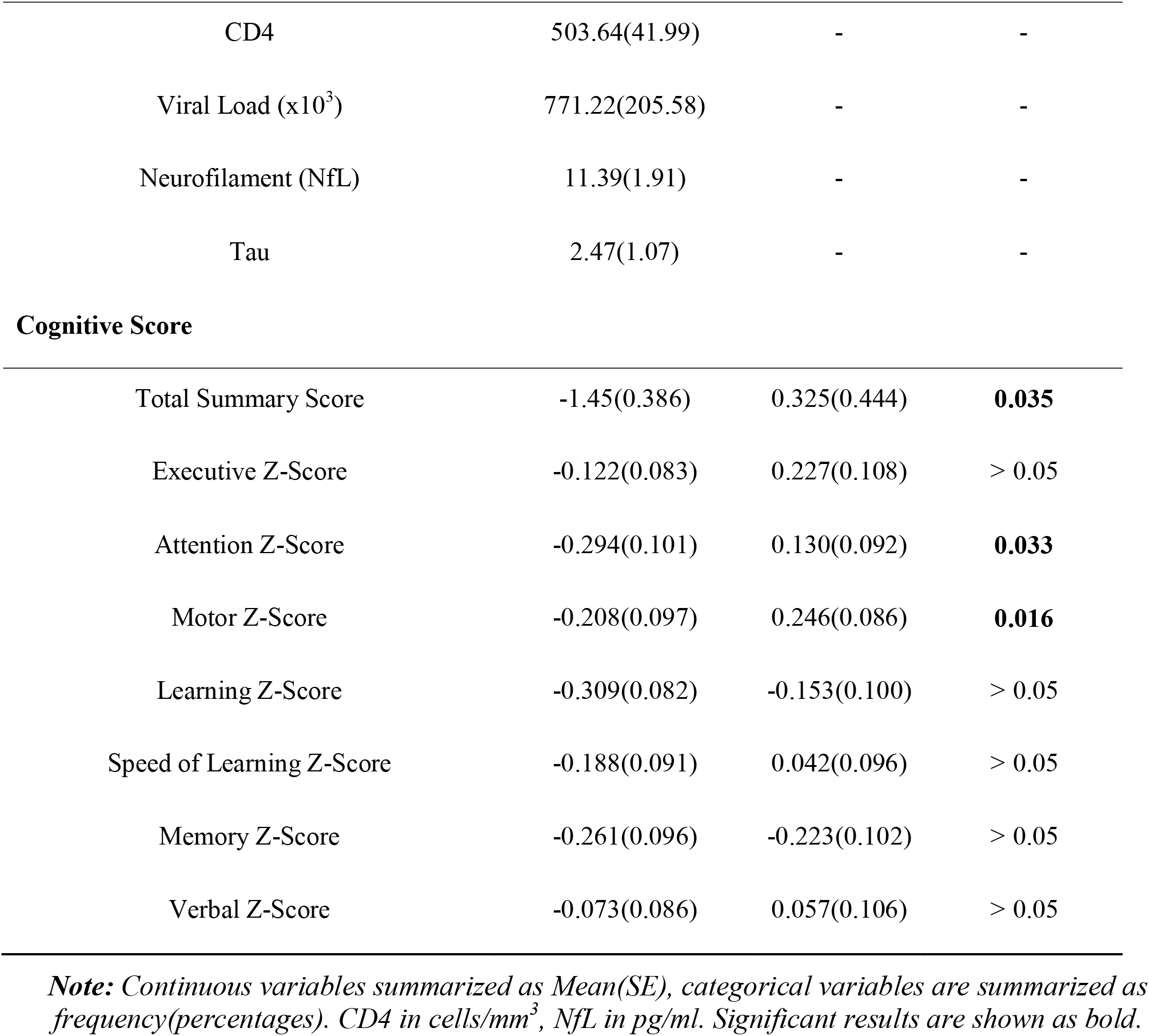
Participant demographics, clinical and cognitive information

|  | **HIV+ (N=42)** | **HIV-(N=52)** | **P-value** |
| --- | --- | --- | --- |
| **Age, mean(SE)** | 34.05(2.01) | 37.13(1.65) | 0.24 |
| **Gender, n(%)** |  |  |  |
| Female | 2(0.048) | 26 (0.5) | <**0.001** |
| Male | 40 (0.952) | 26 (0.5) |  |
| **Ethnicity, n(%)** |  |  |  |
| Hispanic or Latino | 2(0.048) | 3(0.058) | 0.829 |
| Not Hispanic or Latino | 40(0.952) | 49(0.942) |  |
| **Race, n(%)** |  |  |  |
| Caucasian | 20(0.476) | 43(0.827) | < **0.001** |
| Black.A | 21(0.5) | 5(0.096) |  |
| Other | 1(0.024) | 2(0.038) |  |
| Missing | 0(0) | 1(0.019) |  |
| **Education, n(%)** |  |  |  |
| < 12 years | 2(0.048) | 2(0.038) | **0.002** |
| 12 Years (HS Grad) | 11(0.26) | 4(0.077) |  |
| 12-15 Years (Some College) | 16(0.38) | 10(0.19) |  |
| 16 Years (Bachelor's) | 7(0.17) | 21(0.40) |  |
| > 16 Years (Post-Graduate) | 6(0.14) | 15(15.52) |  |
| **Blood Markers, mean(SE)** |  |  |  |
| CD4 | 503.64(41.99) | - | - |
| Viral Load (x10<sup>3</sup>) | 771.22(205.58) | - | - |
| Neurofilament (NfL) | 11.39(1.91) | - | - |
| Tau | 2.47(1.07) | - | - |
| **Cognitive Score** |  |  |  |
| Total Summary Score | -1.45(0.386) | 0.325(0.444) | **0.035** |
| Executive Z-Score | -0.122(0.083) | 0.227(0.108) | > 0.05 |
| Attention Z-Score | -0.294(0.101) | 0.130(0.092) | **0.033** |
| Motor Z-Score | -0.208(0.097) | 0.246(0.086) | **0.016** |
| Learning Z-Score | -0.309(0.082) | -0.153(0.100) | > 0.05 |
| Speed of Learning Z-Score | -0.188(0.091) | 0.042(0.096) | > 0.05 |
| Memory Z-Score | -0.261(0.096) | -0.223(0.102) | > 0.05 |
| Verbal Z-Score | -0.073(0.086) | 0.057(0.106) | > 0.05 |

**Note:** Continuous variables summarized as Mean(SE), categorical variables are summarized as frequency(percentages). CD4 in cells/mm<sup>3</sup>, NfL in pg/ml. Significant results are shown as bold.

### 2.2 Data Acquisition

#### 2.2.1 Blood Sample

Plasma levels of markers associated with neuroinflammation and neurodegeneration (Neurofilament light chain NfL, and Tau protein) were measured by Simoa assay via commercial lab, QuanterixTM (Lexington, MA, United States, *https://www.quanterix.com/).* Viral load (VL) from each HIV+ participant was measured via Roche COBAS 8800 System with a lower limit of detection of 20 copies/mL. CD4+ count was obtained via flow cytometric immunophenotyping at the Clinical Laboratory Improvement Amendments (CLIA), certified clinical lab at the University of Rochester.

#### 2.2.2 Neuropsychological assessments

The neurocognitive evaluation was performed by trained staff and supervised by a clinical neuropsychologist. Tests of Executive Function (Trailmaking Test Parts A & B, Stroop Interference task), Speed of Information Processing (Symbol Digit Modalities Test and Stroop 2 Color Naming), Attention and Working Memory (CalCAP(CRT4) and WAIS-III Letter-Number Sequencing), Learning (Rey Auditory Verbal Learning Test AVLT (trials 1-5), Rey Complex Figure Test Immediate Recall), Memory (Rey Auditory Verbal Learning Test RAVLT Delayed Recall, Rey Complex Figure Test Delayed Recall) and Motor (Grooved Pegboard, left and right hand)were administered at each visit. Premorbid intellectual functioning ability was estimated via WRAT-4 Reading at the baseline visit only. Raw scores were converted to z-scores using test manual norms. Cognitive domain scores were created by averaging the z-scores of tests within each domain. A total summary score was calculated by summing the z-scores of the six cognitive domains measured (Executive Function, Speed of Information Processing, Attention and Working Memory, Learning, Memory, and Motor). HAND diagnoses were determined for each participant according to the Frascati criteria (45). Subjects were accordingly defined as either within normal limits (WNL), or cognitively impaired (CI) [i.e., subjects having asymptomatic neurological impairment (ANI) or minor neurocognitive impairment (MND)].

#### 2.2.3 Image Acquisition

All participants were scanned on a 3T MRI scanner ((MAGNETOM Trio, Siemens, Erlangen, Germany) equipped with a 32-channel head coil.

##### Anatomical Imaging

For the purpose of segmentation and identification of the anatomical landmarks, a T1-weighted (T1w) images were acquired using a 3D magnetization prepared rapid acquisition gradient-echo (MPRAGE) sequence with Inversion Time (TI) = 1,100 ms, Repetition Time (TR) = 2,530 ms, Echo Time (TE) = 3.44 ms, Flip Angle = 7 ◻, Field of View (FOV) = 256×256; GRAPPA factor = 2, number of average =1, number of slices = 192, voxel size = 1.0×1.0×1.0 mm^3^, and total time of acquisition (TA) was 5:52 min.

##### Diffusion Tensor Imaging

Diffusion weighted images (DWI) were acquired using a single shot spin echo echo-planar imaging (SE-EPI) sequence with 60 non-collinear diffusion-encoded images (b =1000 s/mm^2^), 10 non-diffusion weighted reference images (b=0 s/mm^2^); TR = 8,900 ms; TE = 86 ms; FOV = 256×256; GRAPPA factor = 2; number of slices= 70; number of volumes = 61; voxel size = 2.0×2.0×2.0mm^3^; TA = 10:51 min. In order to correct for EPI distortions, a double-echo gradient echo field map sequence was also acquired (TR = 400 ms; TE = 5.19 ms; FOV = 256×256; flip angle = 60 ◻; number of slices = 70; voxel size = 2.0×2.0×2.0 mm^3^; TA = 3:28 min).

#### 2.2.4 Image Preprocessing

All MRI images were visually inspected for any severe artifacts. DWI images were corrected for eddy current-induced distortion, susceptibility-induced distortion, and motion correction using TOPUP and EDDY tools in FSL (*https://fsl.fmrib.ox.ac.uk/fsl/fslwiki/*) (46, 47).

##### 2.2.4.1 Fixel Based Analysis

Fixel-based analysis (FBA) was performed using the recommended pipeline in MRtrix3 (*www.mrtrix.org, version 3.0.2*) (35). Briefly, DWI images were up-sampled by a factor of 2 in all three dimensions using cubic b-spline interpolation. The fiber orientation distributions (FODs) within each voxel were computed using single-shell, 3-tissue constrained spherical deconvolution (SS3T-CSD), using group averaged response functions for WM, GM and CSF (43). A study-specific template was then created by spatial normalization of subjects using symmetric diffeomorphic non-linear transformation FOD-based registration (48). One group-averaged FOD template was created for cross-sectional analysis, including 20 HIV+ and 20 HIV− individuals. The FOD image for each subject was then registered to the template using FOD-guided non-linear registration.

A tractogram was then generated using whole-brain probabilistic tractography on the FOD population template (49). Twenty million streamlines were generated and subsequently filtered to two million streamlines using spherical deconvolution informed filtering of tractograms (SIFT) to reduce reconstruction biases (50). Fixel specific measures of fiber density (FD) and fiber bundle cross-section (FC) were calculated within each voxel. The log of FC (logFC) was calculated to ensure FC values were centered around zero and normally distributed. A combine measure, FDC, was also generated by multiplying FD and logFC. Modified fixel-based metrics were also calculated for the major fiber bundles using the Johns Hopkins University (JHU) DTI-based WM atlas.

##### 2.2.4.2 DTI preprocessing

DTI metrics (FA and MD) were computed using DTIFIT in FSL (51). Free water corrected DTI (fwcDTI) metrics (FA_T_ and MD_T_) were computed with a bi-tensor model from the DWI using a previously described algorithm (52) and the processing was performed using Nextflow pipeline (53) with all software dependencies bundled in a Singularity Container (54).

### 2.3 Statistical Analysis

#### 2.3.1 Subjects Characteristics

Differences in clinical parameters between HIV+ and HIV− cohorts at baseline were examined using two-way independent t-tests at the *α* = 0.05 significance level. Statistical analysis of demographic data was computed in R 3.6.2 (R Foundation for Statistical Computing, Vienna, Austria).

Univariate comparisons between two independent groups were conducted by either two-group Welch’s unequal variances t-test (for continuous variables) or Fisher exact test (for categorical variables). Paired t-tests were used to compare the levels of continuous variables in HIV+ participants between baseline and week-12. Pearson correlation test was used to test the univariate associations between two continuous variables. A p-value < 0.05 was considered statistically significant for a single hypothesis testing problem. For inferential problems that involved multiple hypotheses, Benjamini–Hochberg multiple testing procedure was used to control the false discovery rate (FDR) at *α* < 0.05 level (55).

#### 2.3.2 Fixel-based analysis

##### 2.3.2.1 Whole-brain fixed based analysis

Statistical analyses of images were performed in MRtrix3 (*www.mrtrix.org, version 3.0.2*). All WM fixels were compared between HIV+ and HIV− individuals. Group comparisons were performed for FD, logFC, and FDC at each fixel using a General Linear Model (GLM), with age and sex included as covariates. Connectivity based fixel enhancement (CFE) and non-parametric permutation testing over 5000 permutations were used to identify significant differences in fixel-based metrics (56). Family-wise error (FWE)-corrected p values are reported to account for multiple comparisons. Significant fixels (FWE corrected p-value < 0.05) were visualized using the *mrview* tool in MRtrix3. Fixels were mapped to streamlines of the template-derived tractogram, only displaying streamlines corresponding to significant fixels. Significant streamlines were colored by the effect size, presented as a percentage relative to HIV− individuals or by streamline direction (left-right: red, inferior-superior: blue, anterior-posterior: green).

##### 2.3.2.2 Region of interest analysis

Region of interest (ROI) analysis was performed for fixel-based metrics (FD, logFC and FDC), DTI metrics (FA and MD), fwcDTI metrics (FA_T_ and MD_T_) using the JHU DTI-based WM atlas. The following ROIs were included in the analyses: the left and right posterior limb of internal capsule (PLIC), the left and right superior corona radiata (SCR), the left and right cerebellar peduncles (CP), the left and right inferior cerebellar peduncle (ICP), and the middle cerebellar peduncle (MCP). These regions were chosen *a priori* based on the findings from whole brain fixel-based analysis. The mean value for FD, logFC, and FDC was computed for each ROI and compared across groups. Correlation analyses were also performed to evaluate the relationship between fixel-based metrics and cognitive z-scores. Independent t-tests with multiple comparisons corrections were used to compare mean logFC, FD, and FDC between cohorts over the ROIs. Correlations were performed using the non-parametric Spearman’s rho and a linear model with age and sex as covariates. Benjamini-Hochberg procedure was applied to control the FDR at α < 0.05 significance level.

#### 2.3.3 Tract Based Spatial Statistics (DTI and fwcDTI)

FSL based TBSS was performed to investigate the FA, free water corrected FA (FA_T_), MD and free water corrected MD (FA_T_) changes along WM tracts (24). Group comparisons were performed using FSL Randomize for 5000 permutations. Threshold-free cluster enhancement (TFCE) (57) was used for multiple comparison correction at the α = 0.05 significance level.

#### 2.3.4 Machine Learning Classification

Machine learning classification was performed using FBA and fwcDTI metrics. Classifiers were implemented in scikit-learn (58). Instances were standardized prior to training and dimensionality reduction was performed using kernel principal component analysis (kPCA). Four binary classifiers were used to evaluate the specificity of both fixel based metrics and fwcDTI metrics in their ability to distinguish between WNL and CI. In this study we implemented random forest, naïve bayes, linear discriminant analysis (LDA), and adaptive boosting (AdaBoost) classifiers. All classifiers were optimized using a grid search algorithm with a five-fold cross-validation. Classifiers were evaluated using the weighted average (across classes) for precision, recall and f1-score. Precision, also known as the positive predictive value (PPV), is defined as the number of instances classified as positive, divided by the total number of positive (CI) instances. Recall, or sensitivity, is the number of instances accurately classified as positive (true positives), divided by the total number of instances classified as positive. The F-score is the harmonic mean of precision and recall. Receiver operating characteristic (ROC) curves and precision-recall curves were also evaluated to assess the performance of these classifiers. Results are reported as the average across five-folds.

## 3 Results

### 3.1 Participant Characteristics

Clinical characteristics, demographic information, and cognitive scores for HIV+ and HIV− individuals are presented in **Table 1**. HIV+ and HIV− cohorts did not significantly differ in age, or ethnicity. The total summary z-score, attention z-score, and motor z-score were found to be significantly lower in the HIV+ cohort (p<0.05).

### 3.2 Whole-brain fixel-based analysis

Whole-brain FBA is shown in **Figure 1**. Streamlines corresponding to significant fixels (FWE corrected p-value < 0.05) are represented as the percentage decrease in HIV+ individuals compared to HIV− individuals for FD, logFC, and FDC. Macrostructural decreases (measured via logFC) of up to 15 % were observed along specific fiber tracts. Specifically, the PLIC and MCP were affected bilaterally. The right SCR was also affected. Similar findings were observed for FD, though much less pronounced. Moreover, decreases in FD were more localized to the PLIC. FDC exhibited similar patterns of micro- and macro-structural degeneration, with a larger effect size (**Table 2**). Compared to HIV− individuals, HIV+ individuals had a 35 % decrease in FDC in the PLIC bilaterally as well as the right SCR. **Figure 2** shows streamlines displayed and colored based on orientation for significant decreases in logFC in HIV+ individuals. **Figure 3** shows a coronal view of fiber tract specific significant fixels, and the inset shows zoomed in area indicating regions with crossing fibers around cerebellar peduncles (CP) and MCP.

**Figure 1.**
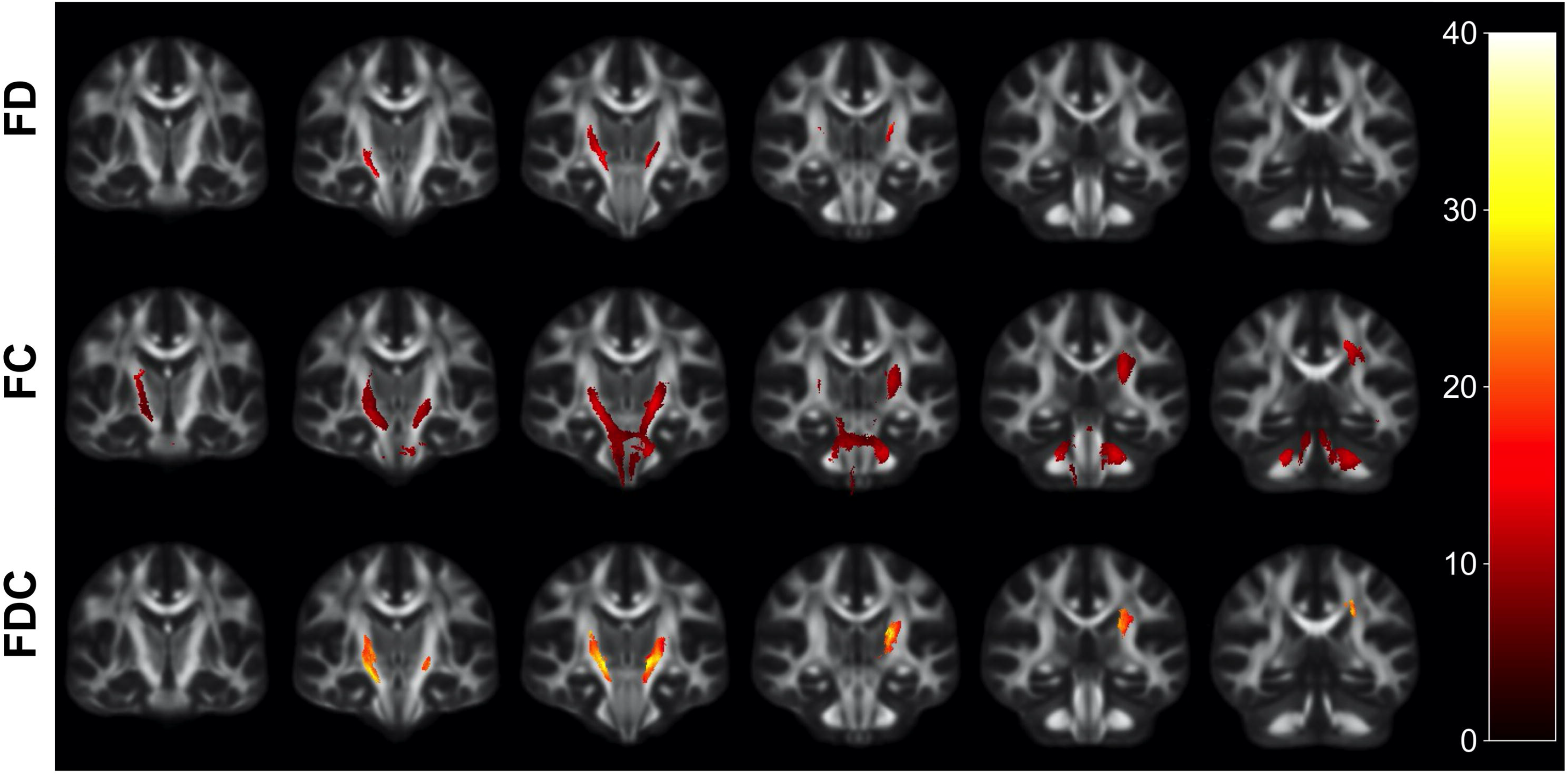
Fiber tract-specific reductions in HIV+ compared to HIV− using whole-brain FBA. Significant fixels (FWE-corrected p-value) between HIV+ and HIV− groups displayed as the percentage decrease in the HIV+ group compared to healthy controls, displayed in coronal slices. FD: fiber density; logFC: fiber bundle cross section; FDC: fiber density and cross-section.

**Table 2.**
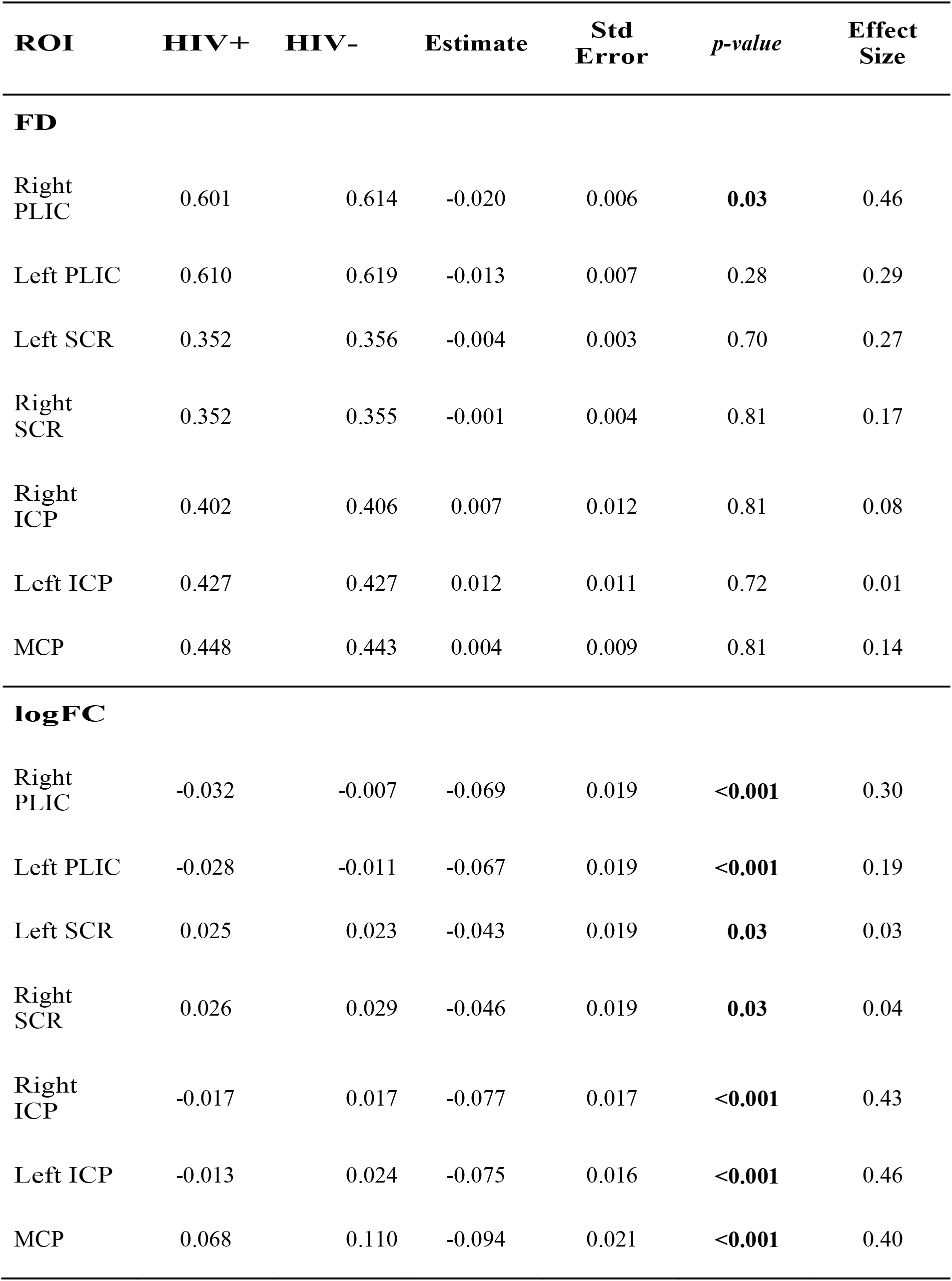
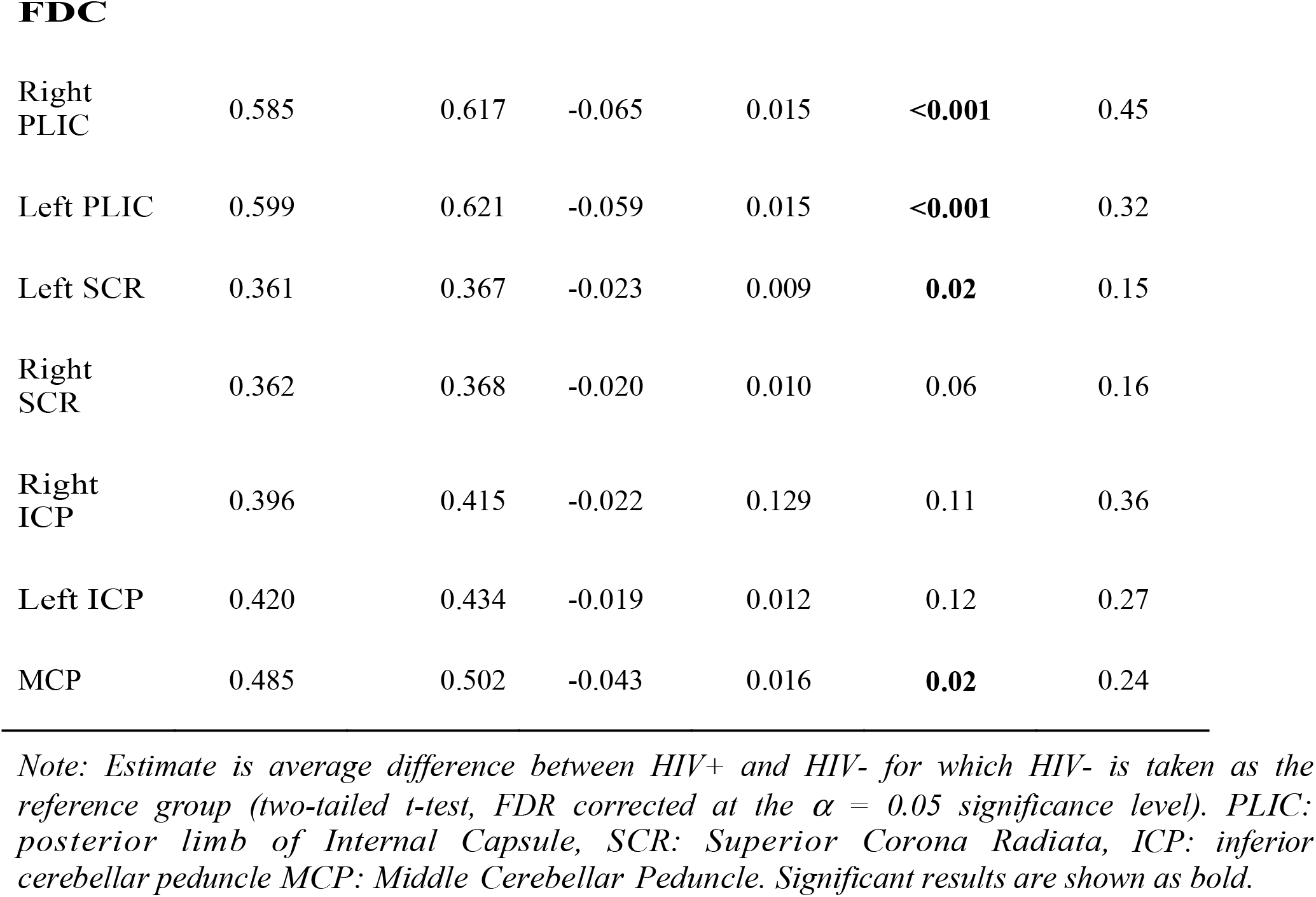
Linear regression model comparing the FBA metrics in HIV+ and HIV− individuals, with age and sex included as covariates.

| **ROI** | **HIV+** | **HIV-** | **Estimate** | **Std Error** | ***p-value*** | **Effect Size** |
| --- | --- | --- | --- | --- | --- | --- |
| **FD** |  |  |  |  |  |  |
| Right PLIC | 0.601 | 0.614 | -0.020 | 0.006 | **0.03** | 0.46 |
| Left PLIC | 0.610 | 0.619 | -0.013 | 0.007 | 0.28 | 0.29 |
| Left SCR | 0.352 | 0.356 | -0.004 | 0.003 | 0.70 | 0.27 |
| Right SCR | 0.352 | 0.355 | -0.001 | 0.004 | 0.81 | 0.17 |
| Right ICP | 0.402 | 0.406 | 0.007 | 0.012 | 0.81 | 0.08 |
| Left ICP | 0.427 | 0.427 | 0.012 | 0.011 | 0.72 | 0.01 |
| MCP | 0.448 | 0.443 | 0.004 | 0.009 | 0.81 | 0.14 |
| **logFC** |  |  |  |  |  |  |
| Right PLIC | -0.032 | -0.007 | -0.069 | 0.019 | **<0.001** | 0.30 |
| Left PLIC | -0.028 | -0.011 | -0.067 | 0.019 | **<0.001** | 0.19 |
| Left SCR | 0.025 | 0.023 | -0.043 | 0.019 | **0.03** | 0.03 |
| Right SCR | 0.026 | 0.029 | -0.046 | 0.019 | **0.03** | 0.04 |
| Right ICP | -0.017 | 0.017 | -0.077 | 0.017 | **<0.001** | 0.43 |
| Left ICP | -0.013 | 0.024 | -0.075 | 0.016 | **<0.001** | 0.46 |
| MCP | 0.068 | 0.110 | -0.094 | 0.021 | **<0.001** | 0.40 |
| FDC |  |  |  |  |  |  |
| Right PLIC | 0.585 | 0.617 | -0.065 | 0.015 | **<0.001** | 0.45 |
| Left PLIC | 0.599 | 0.621 | -0.059 | 0.015 | **<0.001** | 0.32 |
| Left SCR | 0.361 | 0.367 | -0.023 | 0.009 | **0.02** | 0.15 |
| Right SCR | 0.362 | 0.368 | -0.020 | 0.010 | 0.06 | 0.16 |
| Right ICP | 0.396 | 0.415 | -0.022 | 0.129 | 0.11 | 0.36 |
| Left ICP | 0.420 | 0.434 | -0.019 | 0.012 | 0.12 | 0.27 |
| MCP | 0.485 | 0.502 | -0.043 | 0.016 | **0.02** | 0.24 |

*Note: Estimate is average difference between HIV+ and HIV- for which HIV- is taken as the reference group (two-tailed t-test, FDR corrected at the $\alpha = 0.05$ significance level). PLIC: posterior limb of Internal Capsule, SCR: Superior Corona Radiata, ICP: inferior cerebellar peduncle MCP: Middle Cerebellar Peduncle. Significant results are shown as bold.*

**Figure 2.**
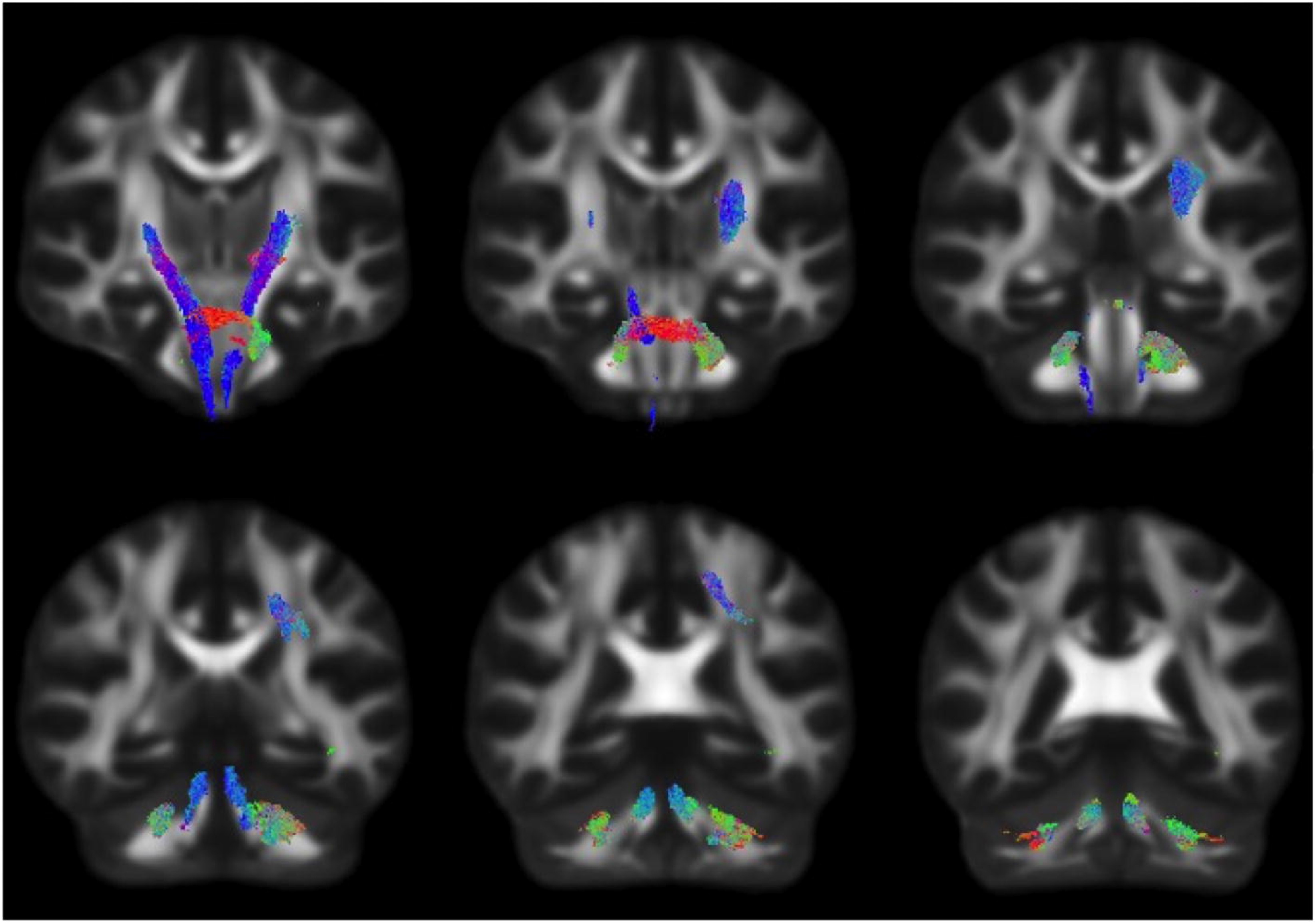
Fiber tract specific logFC decreases in HIV infection, colored by direction. Streamlines were cropped from the template tractogram to include only significant fixels (FWE-corrected p-value < 0.05) for which the logFC metric is decreased in the HIV+ cohort compared to HIV− individuals. Significant streamlines are shown across coronal slices and colored by direction (anterior-posterior: green; superior-inferior: blue; left-right: red).

**Figure 3.**
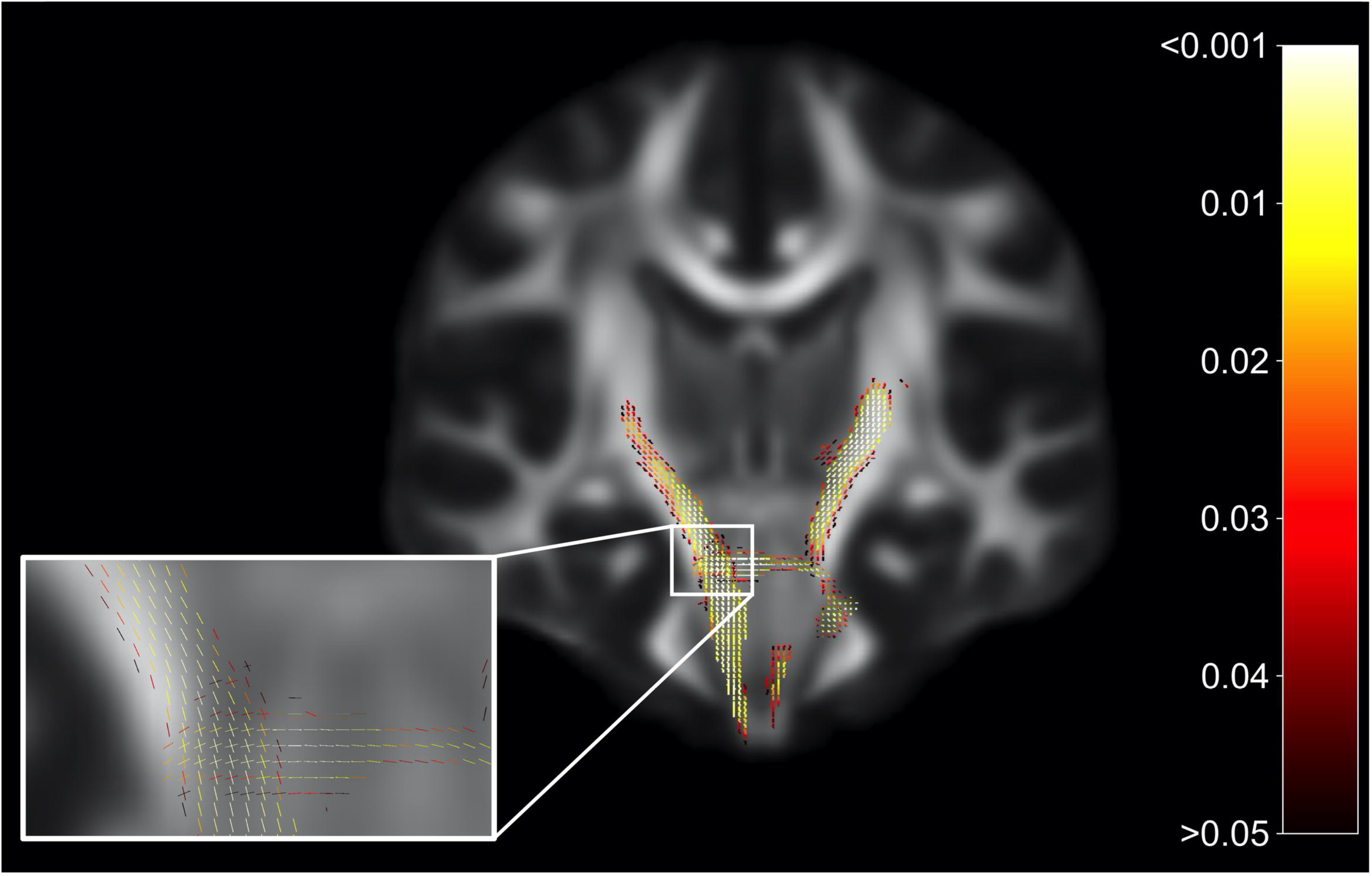
Fiber tract-specific significant fixels. Coronal slice showing fixels that were significantly decreased (FWE-corrected p-values) in HIV+ individuals compared to HIV−. The zoomed in area illustrates differences and p-values assigned to individual fixels in regions with crossing-fibers, around the cerebral peduncles (CP) and middle cerebellar peduncles (MCP). Fixels are colored by FWE corrected p-value.

#### 3.2.1 Region of interest analysis

**Table 2** lists the mean and standard errors for several ROIs between the participants for the FBA metrics FD, logFC, and FDC for HIV+ and HIV− individuals. Linear models were further implemented to evaluate the relationship between fixel-based metrics and cohorts, including age and sex as covariates. We found a significant reduction in several ROIs in FBA metrics in the HIV+ than HIV− individuals. Linear regression models comparing DTI and fwcDTI metrics between HIV+ and HIV− cohorts are provided in **Supplementary Table 1**. None of the ROIs show any significant differences between the cohorts for the DTI and fwcDTI metrics.

**Figure 4(A-C)** shows scatterplots examining the relationship between the attention domain z-scores and FBA metrics (FD, logFC and FDC) for the right and left of PLIC and SCR. For HIV+ individuals, the right PLIC was found to be significant for FD (ρ=0.33, p=0.036), FDC (ρ=0.44, p=0.0039) and logFC (ρ=0.4, p=0.0085) while the right and left SCR were found to be significant in FDC (ρ=0.43, p=0.0048, and ρ=0.38, p=0.014, respectively). On the other hand, none of the ROIs for any metrics were significantly correlated with attention z-scores in the HIV− individuals. **Figure 4D** illustrates the corresponding ROIs.

**Figure 4.**
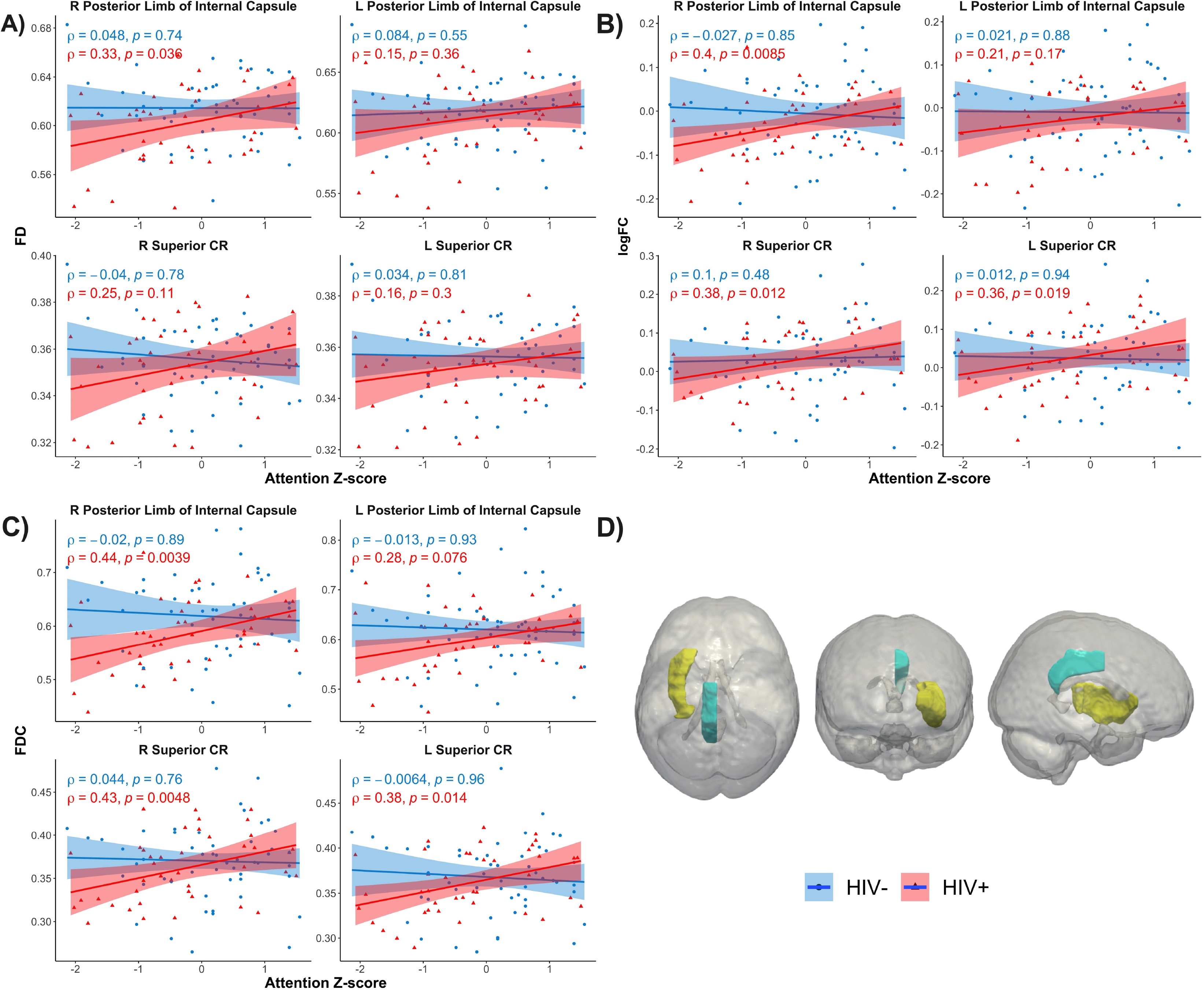
Scatterplots show attention z-score as a function of FBA metrics: A) Fiber Density (FD), B) Fiber Cross-section (logFC), C) Fiber density and cross section (FDC), D) JHU white matter atlas and corresponding regions of interest. Only significant regions shown. Solid lines represent linear fit, and shaded areas represent the 95% confidence interval. IC: internal capsule (yellow); CR: corona radiata (teal); CR: corona radiata.

Additionally, the right PLIC was also found to be significantly correlated with tau protein in HIV+ individuals (ρ=0.32, p=0.043) (Supplementary **Figure S1**). However, no significant associations were observed between FBA metrics and NfL.

### 3.3 Tract Based Spatial Statistics (DTI and fwcDTI)

No significant differences between HIV+ and HIV− cohorts were observed in FA and FA_T_ using TBSS. However, a decrease in FA and FA_T_ (1-2%) was observed in the PLIC, SCR and superior longitudinal fasciculus (SLF) using TBSS (**Figure 5).** TBSS of FA_T_, while not significant, highlighted areas that were decreased in HIV+ individuals compared to HIV− individuals. It should be noted that in our study, the TBSS figures (Figure 5) report regions found for ***p<0.5***. None of the regions survived for significance thresholding for *p<0.05*.

**Figure 5.**
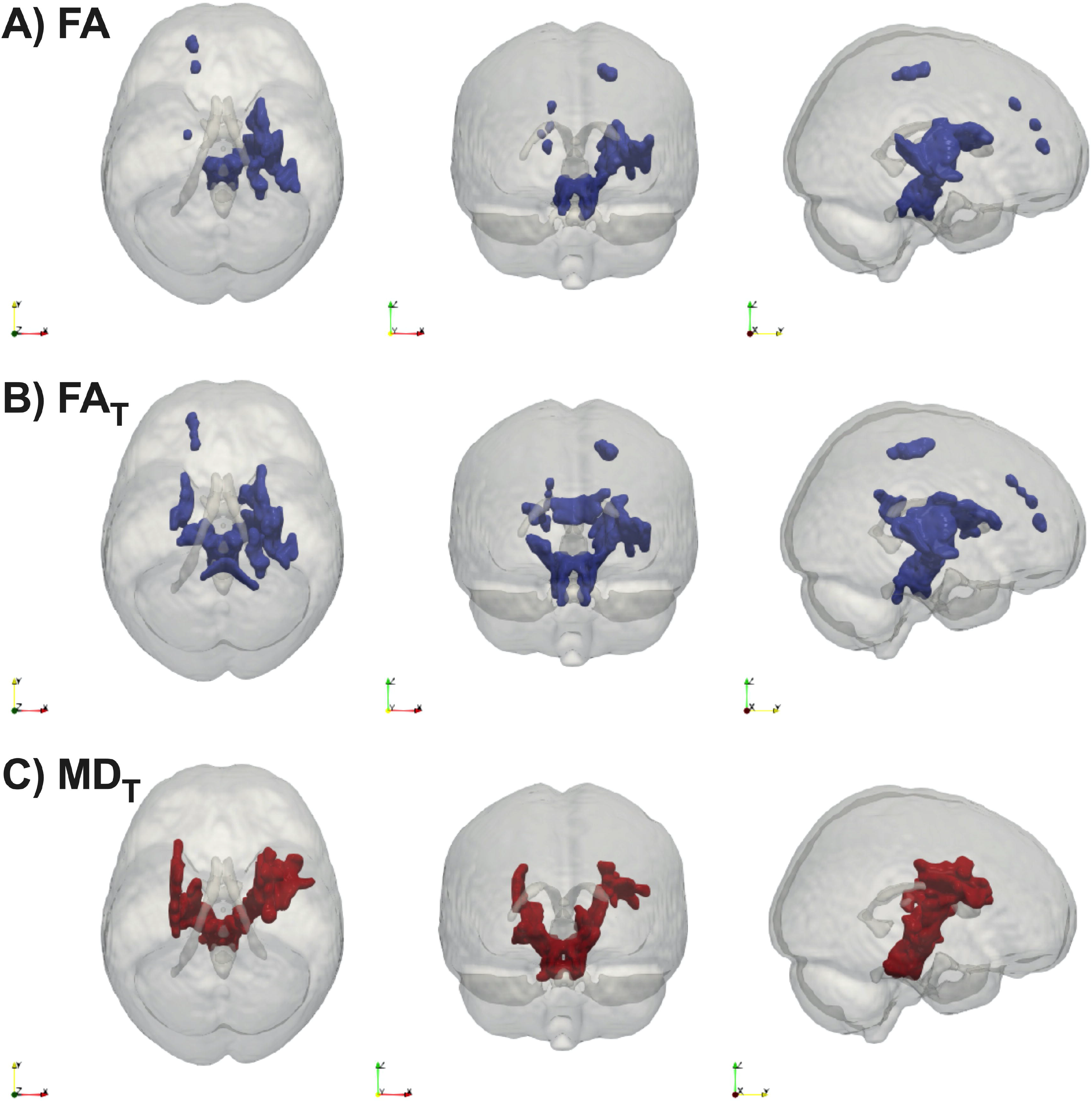
Group comparisons of A) FA, B) FA_T_ C) MD_T_ using TBSS. Statistical maps thresholded at 0.5 (not significant) show regions where FA and FA_T_ are reduced in HIV+ individuals compared to HIV-individuals. FA: Fractional Anisotropy; FA_T_: free water corrected Fractional Anisotropy; Blue represents reduced FA or FA_T_ in the HIV+ compared to HIV− individuals.

### 3.4 Machine Learning Classification

Overall, we found that use of fixel based metrics resulted in a higher precision and recall compared to when using fwcDTI metrics. The adaptive boosting (AdaBoost) method resulted in the best performance for fixel based metrics. **Figure 6** shows the ROC and PRC curves using an AdaBoost multiclass classifier. **Table 3** represents the sample averaged precision, recall and f1-scores for other classifiers contrasting specificity of FBA and fwcDTI using five-fold cross validation.

**Figure 6.**
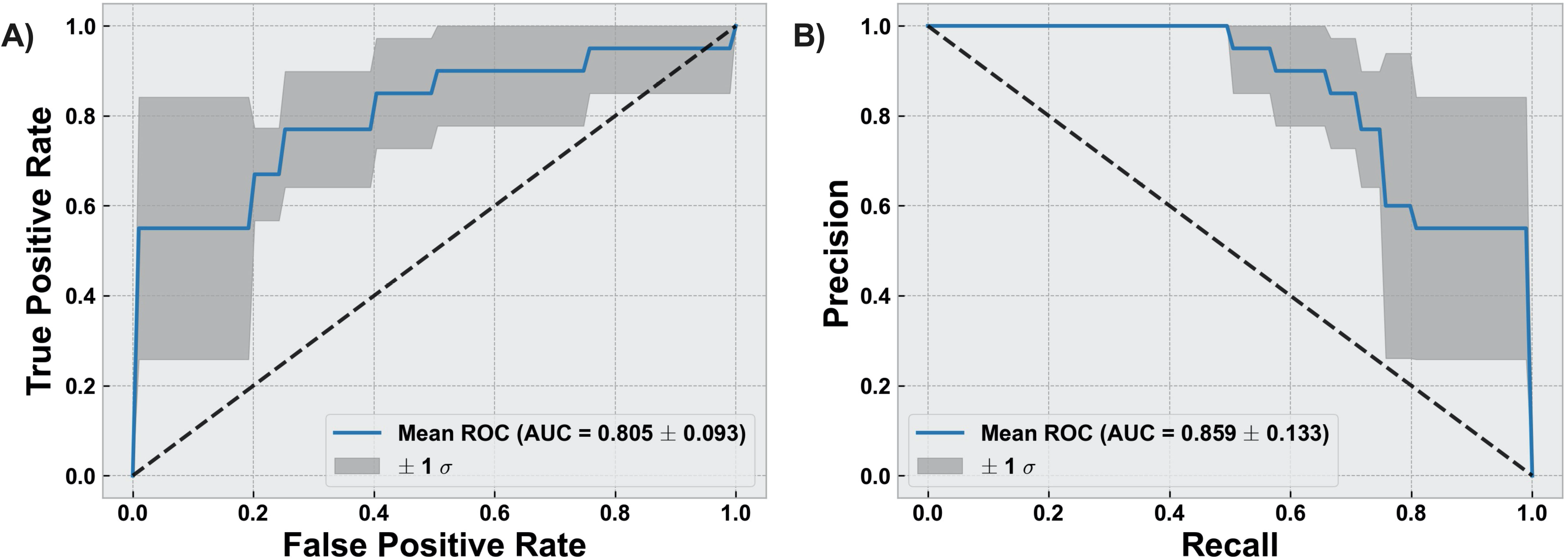
Evaluation of classification results using AdaBoost classifier. **(A)** Receiver operating characteristic (ROC) Curve for cognitively normal compared to cognitively impaired (CI). **(B)** Precision vs recall curve (PRC) for cognitively normal compared to CI. Solid line represents the mean curve using five-fold cross validation. Shaded areas represent +/− 1 standard deviation. AUC reported as mean +/− standard deviation across five-folds.

**Table 3.** Precision, recall, and F1-scores for 5 different classifiers for FBA and fwcDTI metrics reported using five-fold cross validation in HIV+ subjects only.

| **Classifiers** | **Metrics** | **Precision** | **Recall** | **F-Score** |
| --- | --- | --- | --- | --- |
| LDA | FBA | 0.62 | 0.60 | 0.58 |
|  | fwcDTI | 0.50 | 0.43 | 0.46 |
| Random Forest | FBA | 0.62 | 0.77 | 0.76 |
|  | fwcDTI | 0.53 | 0.53 | 0.53 |
| NaiveBayes | FBA | 0.62 | 0.58 | 0.55 |
|  | fwcDTI | 0.38 | 0.38 | 0.38 |
| AdaBoost | FBA | **0.80** | **0.77** | **0.76** |
|  | fwcDTI | 0.48 | 0.46 | 0.47 |

Note: Scores reported as weighted average across 3 classes. FBA: Fixel-based analysis; fwcDTI: free water corrected diffusion tensor imaging; LDA: Linear Discriminant Analysis.

## 4 Discussion

In this study, we evaluated fiber tract-specific WM changes in HIV infection using FBA, DTI and fwcDTI metrics. The major findings of this work were that: a) HIV+ individuals exhibit axonal degradation within the PLIC, CP and SCR as revealed by FBA; b) Similar trends were observed using TBSS of FA and FA_T_. In contrast to FA, FA_T_ showed trends towards more areas that were decreased in HIV+ individuals compared to HIV− individuals. c) FBA metrics in PLIC and SCR exhibit significant positive associations with attention cognitive z-scores in HIV+ individuals; d) Machine learning classifiers for FBA reliably distinguished between cognitively normal patients and those with cognitive impairment in patients with HIV infection.

To the best of our knowledge, this is the first study investigating FBA and TBSS of fwcDTI metrics (FA_T_, and MD_T_) in HIV infected individuals. The work presented here provides a comprehensive and robust framework for evaluating brain injury (and secondary chronic inflammation) in the setting of HIV. Chronic neuroinflammation results in damage to the CNS, alteration of the blood brain barrier (BBB), and chronic edema (59). Changes in whole-brain FBA were found along distinct fiber tracts associated with motor and attention cognitive domains. In particular PLIC, CP, MCP, and SCR were affected and exhibited reduced fiber density and fiber bundle cross-section in HIV+ individuals compared to HIV− individuals. ROI based analysis revealed lower mean fixel-based metrics in the HIV+ cohort compared to the HIV− cohort, consistent with those obtained from the whole-brain FBA results.

Previous work using DTI has shown that FA is decreased in corticospinal tract and that MD is increased in the corticospinal tract bilaterally (23). However, FW contamination results in fitting each voxel with an isotropic tensor, leading to an erroneous conclusion that FA is decreased in the presence of edema (12). Consistent with previous work, our findings suggest that axonal degeneration occurs only in fixels associated with the CST in HIV infection, but accounting for edema and FW contamination. Moreover, FBA is a more robust method to evaluate WM structural integrity compared to TBSS, with or without FW correction. This is likely because constrained spherical deconvolution estimates the response function separately for WM, GM, and extracellular FW, ultimately better modeling the response function of WM than DTI (44).

In addition, our findings are consistent with previous work investigating WM in HIV infection, and with the clinical presentation of HAND (2). However, the present study emphasizes FBA to provide a more robust means to evaluate WM structural integrity independent of partial volume effects and FW contamination. Although we only saw trends using TBSS of fwcDTI, it is reasonable to implement this approach to DTI data. Clinically, HAND is a spectrum of disorders in which patients may present with difficulties in cognition, particularly declines in psychomotor processing, attention, and memory. Of interest, the role of the corona radiata in motor pathways is well established, however recent studies have suggested that the corona radiata is related to attention as well (60). As a major WM intersection, it is possible that damage to the corona radiata, observed in this study, affects both corticospinal fibers as well as association fibers passing through the SCR, contributing to the diffuse cognitive changes seen in HAND, particularly psychomotor slowing. Additionally, the default mode network (DMN) has been implicated in HIV and HAND (9). The DMN is primarily composed of the medial prefrontal cortex, posterior cingulate cortex, precuneus and angular gyrus. Moreover, the DMN, a task-negative network is associated with attention and memory. Thus, it is feasible that degeneration of association fibers passing through the corona radiata and internal capsule disrupt connections to the DMN(61).

We also investigated the relationship between fixel based metrics (i.e., FD, logFC and FDC) and inflammatory blood markers including CD4, VL, and neuronal markers NfL, and tau protein in HIV+ individuals. The PLIC was the only structure significantly correlated with tau protein (ρ=0.32, p=0.043). No other blood markers were significantly correlated with fixel-based metrics. Tau protein is a component of the neurofibrillary tangles most often associated with Alzheimer’s Disease (62). However, increasing evidence suggests that chronic neuroinflammation in the setting of HIV infection predisposes HIV+ individuals to premature neurodegeneration as measured by tau protein (63).

Lastly, several machine learning classifiers were used to classify cognitive status in HIV+ individuals using fixel based and free water corrected DTI metrics. In general, we observed that fixel based metrics results in improved performance as measured with precision, recall and f1-score, compared to fwcDTI metrics. Inclusion of other relevant imaging metrics and biomarkers is likely to further improve prediction of developing HAND.

This study has some limitations. First, only 6 of the subjects in the study had mild neurological disorder (MND), therefore our study was mostly composed of cognitively normal subjects and patients with ANI. In this work, MND and ANI were combined and categorized as CI. Second, the proportion of male and female subjects was not equal in HIV+ cohort. However, FBA, and DTI metrics were not significantly different in males vs. females in our HIV− participants, which had a more equal representation. Future research will investigate the utility of fixel based metrics in evaluating HIV associated neuroinflammation longitudinally and developing prognostic machine learning models for predicting HAND longitudinally.

## 5 Conclusions

Our findings suggest that FBA provides a comprehensive and accurate assessment of WM structural integrity in the setting of chronic neuroinflammation in HIV population. Our results indicate that degeneration occurs along specific fiber tracts, which manifests both as macrostructural and microstructural alterations; particularly in the internal capsule and corona radiata in HIV+ individuals. Moreover, our findings are consistent with the clinical presentation of HAND, which often presents as psychomotor slowing with impaired attention, memory, and fine motor function. TBSS based analysis of free water corrected and uncorrected DTI metrics showed decreasing trends between the HIV+ and control group. However, these were not significant, suggesting lower sensitivity for the level of pathology in the cohort under investigation compared to FBA. Therefore, FBA may provide a sensitive biomarker to monitor axonal degeneration in individuals with HIV infection.

## Abbreviations

cART: Combined antiretroviral therapy
CNS: central nervous system
BBB: blood brain barrier
WM: white matter
HAND: HIV associated neurocognitive disorder
FW: free water
DTI: diffusion tensor imaging
FA: fractional anisotropy
AD: axial diffusivity
RD: radial diffusivity
MD: mean diffusivity
TBSS: Tract-based spatial statistics
CSF: cerebrospinal fluid
FA_T_: free water corrected FA
MD_T_: free water corrected MD
FBA: fixel based analysis
FD: fiber density
FC: fiber bundle cross section
FDC: fiber density cross section
AFD: apparent fiber density
FOD: fiber orientation distribution
MT-CSD: multi tissue constrained spherical deconvolution
GM: grey matter
NfL: neurofilament light chain
VVL: viral load
GLM: general linear model
CFE: connectivity based fixel enhancement
FDR: false discovery rate
ROI: region of interest
TFCE: threshold-free cluster enhancement
WNL: within normal limits
ANI: asymptomatic neurological impairment
MND: minor neurocognitive impairment
CI: cognitively impaired
kPCA: kernel principal component analysis
LDA: linear discriminant analysis
PPV: positive predictive value
ROC: receiver operating characteristic
AUC: area under the curve
fwcDTI: free water corrected DTI
DWI: diffusion weighted imaging
PLIC: posterior limb of the internal capsule
MCP: middle cerebellar peduncles
SCR: superior corona radiata
CP: cerebellar peduncle
SLF: superior longitudinal fasciculus
AdaBoost: adaptive boosting
PRC: precision recall curve
DMN: default mode network
CST: corticospinal tract.

## 6 Acknowledgments

This work was supported by the National Institutes of Health (NIH) grants R01-MH099921, R01-AG054328, and R01-MH118020. We thank our study participants and staff who were involved in image acquisition and clinical/neurocognitive assessments.

## 8 Author Contributions

AF (Alan Finkelstein): Study concept, image processing, data analysis and interpretation, manuscript writing, original draft; AF (Abrar Faiyaz): Image processing, data analysis and interpretation, and manuscript review for intellectual content; MTW: Cognitive data collection and manuscript review for intellectual content; XQ: Statistical interpretation, and manuscript review for intellectual content; MNU: Study concept and interpretation, manuscript writing and review for intellectual content; JZ: Manuscript review for intellectual content; GS: Project administration, funding acquisition, interpretation, manuscript review for intellectual content. All authors contributed to the article and approved the submitted version.

## 9 Data Availability Statement

Anonymized data will be made available on reasonable request, pending appropriate institutional review board approvals.

## 10 Approval for human experiments

All protocols were reviewed and approved by the Research Subjects Review Board (RSRB) at the University of Rochester. All participants signed a written consent form and all experiments were performed in accordance with relevant guidelines and regulations.

