## Supplementary Materials for "Fixel-Based Analysis and Free Water Corrected DTI Evaluation of HIV Associated Neurocognitive Disorders"

**Supplementary Material**

**Supplementary Table S1.** Linear regression model comparing the DTI and fwcDTI metrics in HIV+ and HIV- individuals, with age and sex included as covariates.

| **ROI** | **HIV+** | **HIV-** | **Estimate** | **Std Error** | ***p-value*** | **Effect Size** |
| --- | --- | --- | --- | --- | --- | --- |
| **FA** |  |  |  |  |  |  |
| Right PLIC | 0.590 | 0.592 | -0.0054 | 0.0046 | 0.608 | 0.106 |
| Left PLIC | 0.585 | 0.585 | -0.0038 | 0.0049 | 0.608 | 0.021 |
| Left SCR | 0.442 | 0.439 | 0.0044 | 0.0049 | 0.608 | 0.144 |
| Right SCR | 0.442 | 0.439 | 0.0026 | 0.0049 | 0.608 | 0.134 |
| Right ICP | 0.394 | 0.403 | -0.0139 | 0.0080 | 0.597 | 0.266 |
| Left ICP | 0.381 | 0.386 | -0.0098 | 0.0091 | 0.608 | 0.119 |
| MCP | 0.420 | 0.413 | 0.0038 | 0.0065 | 0.608 | 0.257 |
| **MD** |  |  |  |  |  |  |
| Right PLIC | 7.00e-4 | 7.00e-4 | 3.30e-6 | 3.70e-6 | 0.590 | 0.231 |
| Left PLIC | 7.10e-4 | 7.00e-4 | 1.50e-6 | 3.70e-6 | 0.801 | 0.208 |
| Left SCR | 7.30e-4 | 7.20e-4 | 4.80e-6 | 6.00e-6 | 0.590 | 0.264 |
| Right SCR | 7.10e-4 | 7.10e-4 | 5.90e-6 | 5.90e-6 | 0.590 | 0.261 |
| Right ICP | 8.30e-4 | 8.20e-4 | 2.62e-5 | 1.48e-5 | 0.562 | 0.259 |
| Left ICP | 8.30e-4 | 8.30e-4 | 5.99e-5 | 1.41e-5 | 0.971 | 0.099 |
| MCP | 8.10e-4 | 8.20e-4 | -1.20e-5 | 1.23e-5 | 0.590 | 0.077 |
| **FA_T_** |  |  |  |  |  |  |
| Right PLIC | 0.593 | 0.596 | -0.0061 | 0.0048 | 0.357 | 0.152 |
| Left PLIC | 0.588 | 0.589 | -0.0048 | 0.0052 | 0.502 | 0.040 |
| Left SCR | 0.457 | 0.454 | 0.0010 | 0.0052 | 0.843 | 0.131 |
| Right SCR | 0.454 | 0.452 | 0.0030 | 0.0053 | 0.663 | 0.106 |
| Right ICP | 0.448 | 0.458 | -0.0156 | 0.0073 | 0.246 | 0.293 |
| Left ICP | 0.448 | 0.452 | -0.0099 | 0.0076 | 0.357 | 0.122 |
| MCP | 0.496 | 0.496 | -0.0085 | 0.0054 | 0.357 | 0.013 |
| **MD_T_** |  |  |  |  |  |  |
| Right PLIC | 7.00e-4 | 7.00e-4 | 4.00e-6 | 3.80e-6 | 0.698 | 0.288 |
| Left PLIC | 7.10e-4 | 7.00e-4 | 2.00e-6 | 3.70e-6 | 0.698 | 0.238 |
| Left SCR | 7.20e-4 | 7.20e-4 | 3.60e-6 | 5.60e-6 | 0.698 | 0.260 |
| Right SCR | 7.10e-4 | 7.10e-4 | 5.80e-6 | 5.70e-6 | 0.698 | 0.269 |
| Right ICP | 7.70e-4 | 7.70e-4 | 6.20e-6 | 1.17e-5 | 0.698 | 0.017 |
| Left ICP | 7.80e-4 | 7.90e-4 | 6.40e-6 | 2.89e-5 | 0.826 | 0.063 |
| MCP | 7.30e-4 | 7.60e-4 | -3.94e-5 | 1.64e-5 | 0.130 | 0.408 |

Note: Estimate is average difference between HIV+ and HIV- for which HIV- is taken as the reference group (two-tailed t-test, FDR corrected at the α = 0.05 significance level). PLIC: posterior limb of Internal Capsule, SCR: Superior Corona Radiata, ICP: inferior cerebellar peduncle MCP: Middle Cerebellar Peduncle, fwcDTI: free water corrected DTI.

**Supplementary Figure S1:** Boxplots of cognitive domain Z-scores by HIV-status. Worse scores (negative Z scores) for attention, memory and overall cognitive summary score were observed in the HIV+ individuals compared to HIV uninfected individuals. Significant group differences are shown with p<0.05.

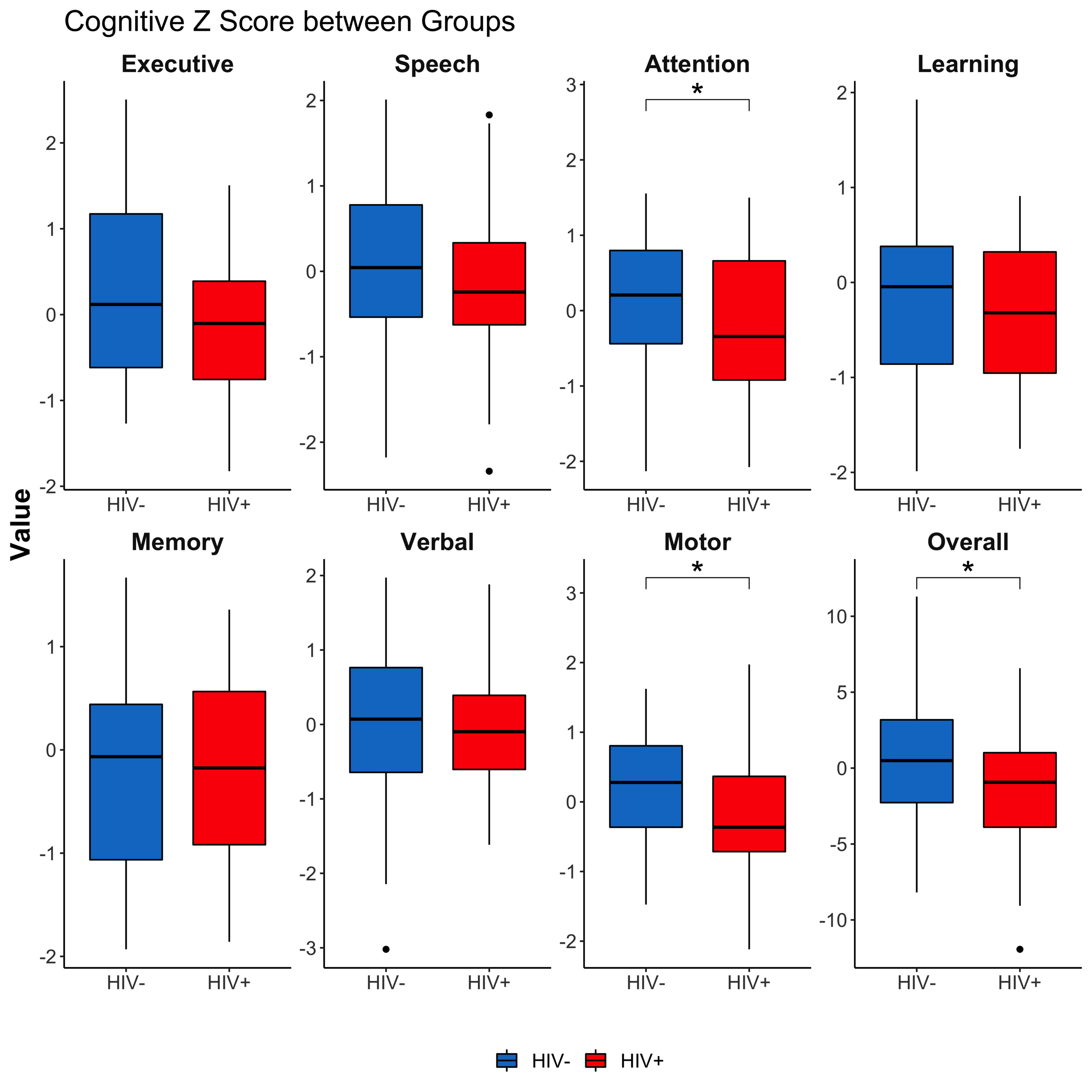

**Supplementary Figure S2:** Relationship between fiber density and cross-section (FDC) and Tau protein in HIV+ cohort. Only significant regions shown. Solid lines represent linear fit, and shaded areas represent the 95% confidence interval. IC: internal capsule.

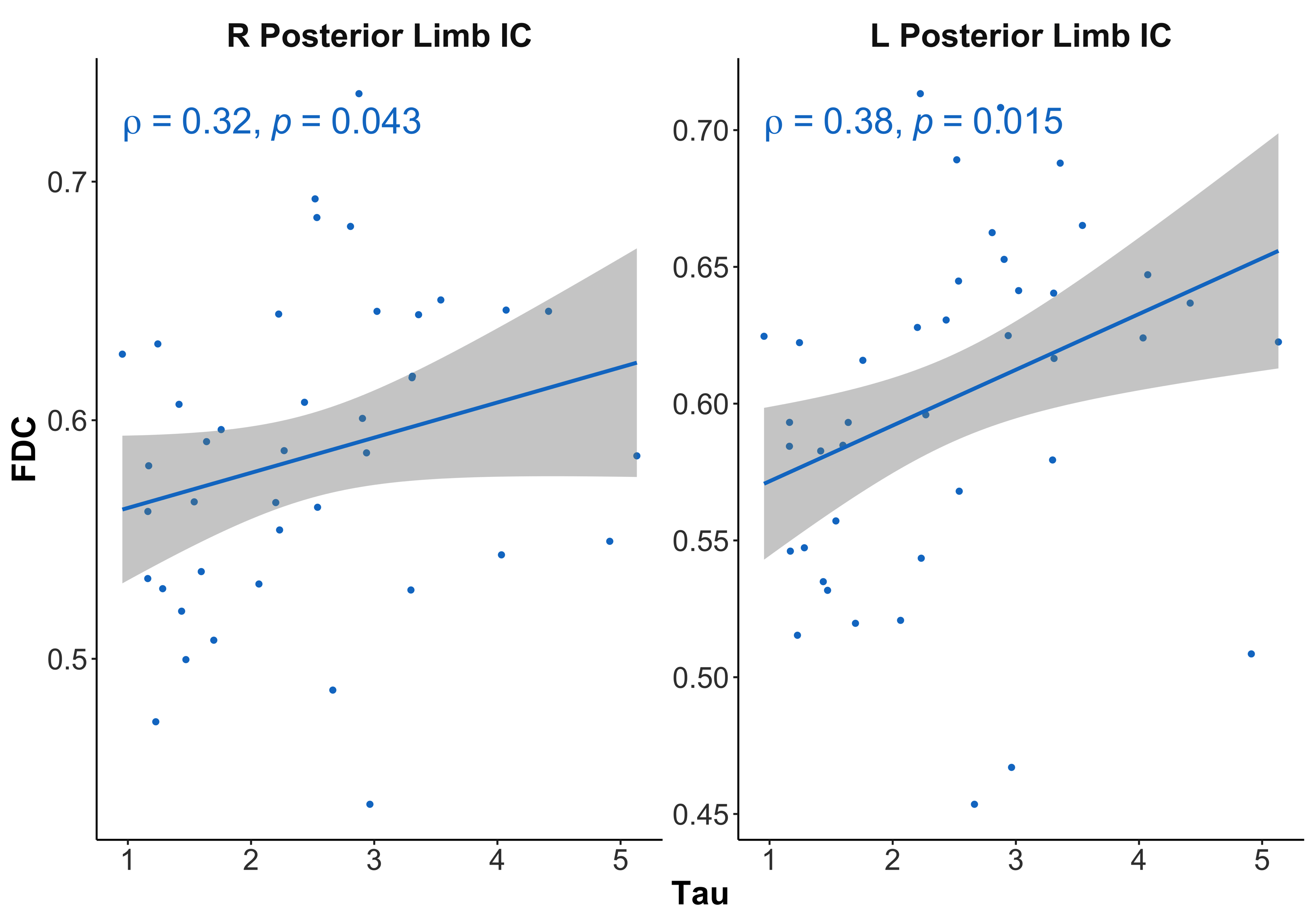

We found that higher levels of tau protein were associated with higher FDC values. One possible explanation is that higher levels of tau in the blood suggest that more tau protein is being cleared from the brain. However, future studies are needed to evaluate levels of tau protein in the CSF and blood to confirm this.
